# Cytosolic delivery of large supramolecular protein complexes arranged on DNA nanopegboards

**DOI:** 10.1101/236729

**Authors:** Pavel M. Nikolov, Katja J. Koßmann, Alessa Schilling, Alessandro Angelin, Josipa Brglez, Alina Klein, Robert Tampé, Kersten S. Rabe, Christof M. Niemeyer

## Abstract

A generic methodology for cytosolic delivery of large supramolecular multiprotein complexes into living cells is described that takes advantage of the highly-controllable bottom-up fabrication of protein-decorated DNA nanostructures and the microfluidic “cell squeezing” technique. Therein, cells are deformed upon passage through a narrow constriction leading to formation of transient holes in the cell membrane that enable the diffusion of the protein-DNA nanostructures from the surrounding buffer into the cytosol. A diverse set of multiprotein complexes was assembled on DNA origami nanostructures using streptavidin and the sensitive glucose sensor protein FLIP as model systems. We demonstrate that our approach allows for the direct cytosolic delivery of these multifunctional protein complexes into the cytosol of HeLa cells. We also demonstrate that targeting groups can be incorporated into the protein-DNA nanoassemblies to enable their intracellular targeting to cytosolic compartments, such as the cytoskeleton or nucleus. We believe that this methodology will open up novel strategies for research in fundamental cell biology, such as the reverse-engineering of the supramolecular machinery involved in gene regulation, cell signalling, or cell division. Furthermore, direct applications in immunotherapy can be foreseen.

---

Engineering of intracellular protein and nucleic acid delivery has been a major challenge in biomedical research for decades^1^, and numerous strategies have evolved to promote, in particular, the cytosolic delivery of biomacromolecular drugs^2^. While recent advances have led to solutions for particular macromolecular species, such as recombinant antibodies tagged with designed endosomolytic or cell-penetrating peptides^3, 4^, generic methodologies for cytosolic delivery of arbitrary proteins or even larger supramolecular aggregates thereof are not known to date^5^. This is unfortunate since the efficient delivery of macromolecules to the cytosol would open the door to novel strategies of research in fundamental cell biology and therapeutics development. For example, cytosolic delivery of large supramolecular protein assemblies could be used in a reverse-engineering approach to elucidate the molecular machinery that is key in all fundamental processes of life, such as gene regulation, cell signaling, or cell division. Because these large complexes self-assemble – often transiently - from tens to hundreds of individual components, their detailed structure, dynamics and function are difficult to analyse. In principle, this can be achieved by reverse-engineering parts of the complexes in order to probe their interactions with distinctive binding partners *in vitro*^6^. However, efficient methodologies for the construction of artificial multiprotein assemblies and their intracellular delivery remain a major challenge for this approach.

DNA nanotechnology arguably offers the most effective means to the rational construction of multiprotein assemblies^7^. The “scaffolded DNA origami” technique^8^ employs long single-stranded DNA (ssDNA) molecules that are folded into arbitrary shapes by aid of short synthetic “staple strand” oligonucleotides. This technique gives ready access to an unlimited variety of finite DNA nanostructures^9, 10^, which can be used as molecular pegboards for the arrangement of non-nucleic acid components, such as proteins, with a precise control over stoichiometry and nanoscale distances^11^. While the resulting multiprotein assemblies are increasingly used as models for the *in vitro* study of spatially-interactive biomolecular networks^12^, such as multienzyme cascades^13^, their exploitation as tools for life sciences remains underdeveloped^14^. Protein-decorated DNA origami nanostructures (DON) have been used *in vitro* to study the coordinated action of motor proteins^15^, they can be used as stable reagents under cell culturing conditions^16–18^, they are functional as immune-activating adjuvants^19, 20^ and drug delivery vehicles^21–24^, and they have been used to bind from the outside of the cell to trigger transmembrane receptors and stimulate signaling^25–28^.

Intracellular delivery of DNA nanostructures is usually achieved via receptor-mediated endocytosis that leads to trafficking of the nanostructures along the cell’s endo-lysosomal pathway^29^. However, the delivery of functional multiprotein-decorated DONs has not yet been realized in this way. Presumably, the harsh environmental conditions inside the endosomal compartments may induce degradation and thus loss of function of the delicate supramolecular architectures. Hence, there is a clear demand for effective methodologies that allow the direct delivery of multiprotein-decorated DONs to the cytosolic compartment of living cells. The method should preserve the DON’s structural integrity and the protein’s tertiary structure, both of which are mandatory to warrant functionality and enable targeting of the protein assemblies to locations of interest, such as the cytoskeleton or nucleus.

Towards this goal, we here demonstrate the utility of a microfluidic method for direct delivery of DNA origami structures into the cytosol of living cells. The microfluidic “cell squeezing” technique, recently developed by the labs of Jensen and Langer^5^, mechanically deforms cells upon passage through a constriction that is smaller than the cell diameter (Figure 1a). The associated compression and shear forces lead to the formation of transient holes in the cell membrane that enable the diffusion of material from the surrounding buffer into the cytosol. Since the method was described as less detrimental than conventional electroporation and highly efficient in the delivery of siRNA, proteins, dextran or even carbon nanotubes,^5^ as well as small molecules^30^, we explored whether microfluidic cell squeezing might be suitable to deliver intact multiprotein-decorated DONs into the cytosolic compartment of living cells. Using a series of increasingly complex protein-decorated DON architectures, we demonstrate that cell squeezing indeed allows for the direct cytosolic delivery and intracellular targeting of supramolecular multifunctional protein-DON assemblies.

**Figure 1.**
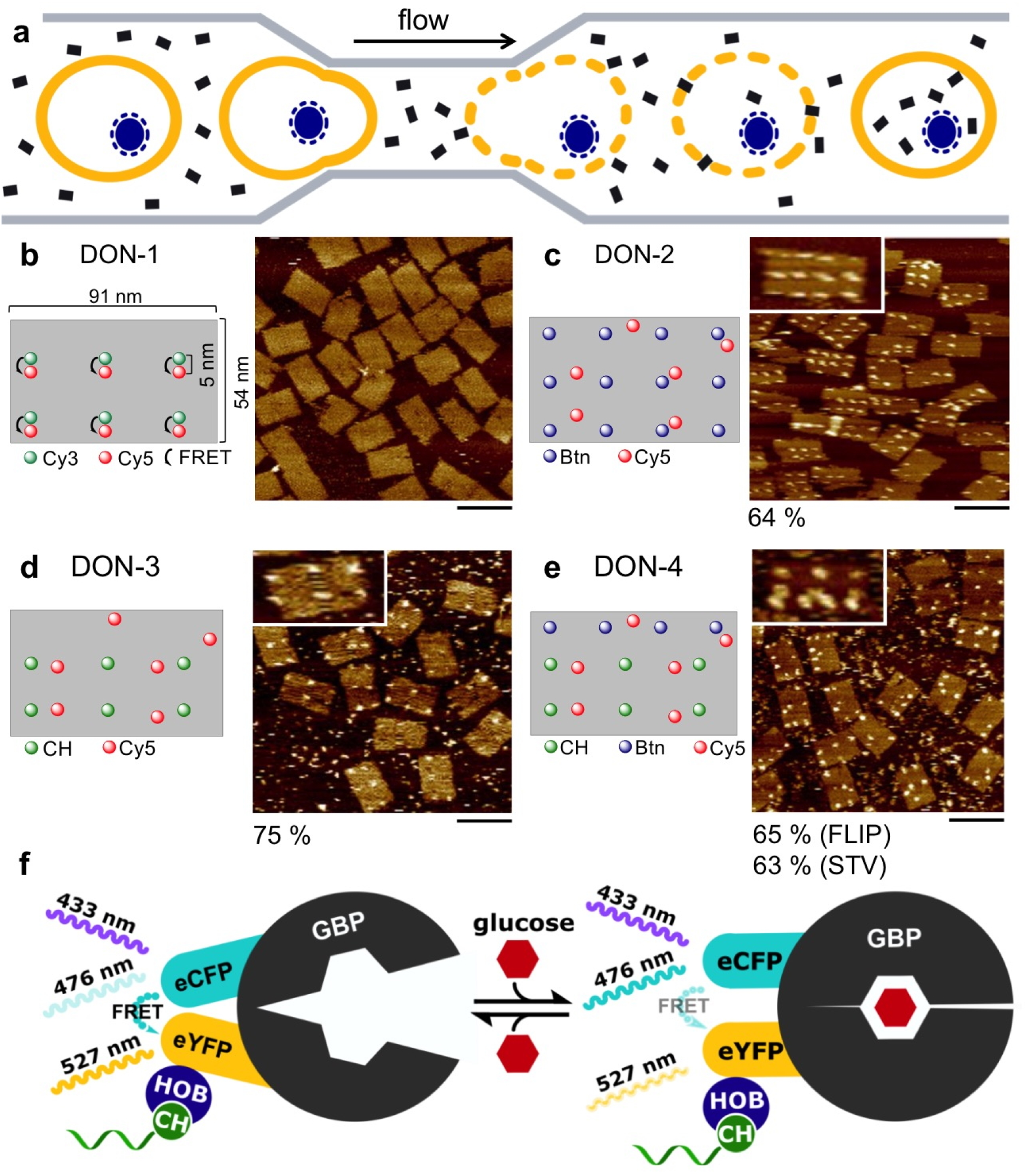
Protein-functionalized DNA origami nanostructures (DON) used for cytosolic delivery by microfluidic cell squeezing. **(a)** Principle of microfluidic cell squeezing. A cell suspension containing the DONs (black rectangles) is squeezed through a microchannel constriction smaller than the cell diameter. Transient formation of pores in the cell membrane enables diffusion of the DONs into the cytosol. Schematic illustrations and representative AFM images of **(b) DON-1** containing 6 Cy3/Cy5 FRET pairs; **(c) DON-2** containing 12 biotin (Btn) and 6 Cy5 modifications for binding of up to 12 streptavidin (STV) proteins; **(d) DON-3** containing 6 chlorohexyl (CH) and 6 Cy5 modifications for binding of up 6 HOB-tagged FLIP proteins; **(d) DON-4** containing 6 CH, 4 Btn and 6 Cy5 modifications for binding of up to six molecules of HOB-tagged FLIP and four STV protein molecules. The percentage of occupied protein binding sites is given underneath the AFM images. Scale bars are 100 nm. (**f**) Schematic drawing of the glucose biosensor FLIP-HOB. Binding of glucose leads to the decrease in FRET and a concomitant change in the fluorescence intensity ratio I(527 nm)/I(476 nm). The HOB domain is used for ligation with chlorohexyl (CH)-modified DNA molecules.

## Results

### Construction of multiprotein-DON assemblies

We assembled the rectangular ~54 × ~91 nm^2^ **DON-1** using the single-stranded 5438 nt template 109Z5, as previously described^31^. **DON-1** contained six Cy3 (donor)/Cy5 (acceptor) Förster resonance energy transfer (FRET) pairs that were incorporated via end-labelled staple strands (Figure 1b) to enable spectroscopic verification of the DNA nanostructure’s integrity in solution. **DON-1** was characterized by AFM analysis on mica surfaces (Figure 1b), gel electrophoresis and fluorescence spectroscopy (Figure S1, Supplementary Information). To investigate whether protein-decorated DONs can be squeezed into cells and targeted to a distinctive cell compartment, we assembled **DON-2** (Figure 1c, Supplementary Figure S2), which contained six Cy5 molecules in addition to twelve biotin groups to facilitate binding of fluorescently labelled streptavidin (STV) or to attach biotinylated targeting groups via STV bridges to the DON. Following this approach, we assembled **DON-2-NLS** (Supplementary Figure S2) by using the biotinylated nuclear localization signal (NLS) peptide (PKKKRKVEDPYC), which is known to carry DNA and other cargo to the cell nucleus^32, 33^. Furthermore, to prove that the squeezing of DNA nanopegboards can be used to deliver functional multiprotein constructs, we assembled **DON-3** that contained six Cy5 labels in addition to six chlorohexane (CH) binding sites (Figure 1d). The CH groups were used as suicide ligands to facilitate covalent binding of self-ligating Halo-tagged fusion proteins^34–36^.

To challenge the concept of squeezing multiprotein nanopegboards, we assembled **DON-3-FLIP** that contained six Cy5 labels and up to six copies of the highly delicate glucose biosensor FLIPglu600μΔ13.^37^ The FLIP protein is a heterotrimeric fusion consisting of glucose binding protein (GBP, colored grey in Figure 1f), which is genetically fused with an enhanced cyan fluorescent protein (eCFP, blue) and an enhanced yellow fluorescent protein (eYFP, yellow) at the N- and C-terminus, respectively. The two fluorescent proteins (FPs) constitute a FRET pair, which is in close proximity in the absence of glucose to facilitate an efficient FRET from eCFP (λ_em, max_ = 476 nm) to eYFP (λ_em, max_ = 527 nm). Upon binding of glucose, GBP adopts a conformation into a closed form, which separates the FRET pair, thereby decreasing the efficiency of energy transfer. As a result, a gradual decrease in fluorescence ratio (*I*(527 nm)/(476 nm)) is observable with increasing glucose concentrations (Supplementary Figure S3). Since the FRET ratio is a highly sensitive measure for even slight changes in secondary and tertiary structure of FLIP, this protein is ideal for detecting any adverse effects on its structure and activity that may result from either the integration into supramolecular DON architecture or the micromechanical transfection procedure. For site-specific immobilization onto DONs, FLIP was genetically fused with a self-labelling Halo-based oligonucleotide binder (HOB) tag that enables covalent ligation with the incorporated CH groups of **DON-3**, in a similar fashion as the regular HaloTag, however, with significantly higher efficacy^38^. Successful loading of **DON-3** with FLIP-HOB was verified by AFM (Figure 1d, see also Supplementary Figure S4). *In vitro* testing revealed that the **DON-3-FLIP** construct functions as a glucose sensor in a similar fashion as the free FLIP-HOB protein (Supplementary Figure S3).

In a modular approach, we used orthogonal DON functionalization and assembled multiprotein constructs bearing FLIP-HOB as a functional domain along with STV to tether Phalloidin (PL) or NLS as targeting domains to the supramolecular constructs. PL is a bicyclic heptapeptide toxin, which strongly binds to filamentous actin (f-actin) and its derivatives containing fluorescent dyes or biotin are routinely used for imaging f-actin in eukaryotic cells^39^. To study the targeting of protein-DON complexes to distinctive locations inside the cytosolic compartment, **DON-4** containing six Cy5 labels as well as six CH and four biotin groups (Figure 1e) was assembled and subsequently modified with FLIP-HOB, streptavidin and biotinylated PL or NLS. The resulting constructs (**DON-4-FLIP-PL, DON-4-FLIP-NLS**) were characterized by AFM (Figure 1e) and gel electrophoresis (Supplementary Figure S5).

### Micromechanical transfection does not disturb the supramolecular integrity

A suspension of HeLa cells in **DON-1** solution (10 nM final concentration) was pressed through the microfluidic squeezing channels. Subsequently, the cells were collected by centrifugation and medium was added to enable cell adhesion and growth. In control experiments, cells were simply cultured for 12 h in the presence of **DON-1** to enable the uptake by endocytosis. Analysis of the cells by confocal fluorescence microscopy (Figure 2a, b) clearly revealed the presence of **DON-1** inside the HeLa cells, as indicated from fluorescence of the Cy3 and Cy5 labels of **DON-1**. Comparison of the micrographs suggested that delivery by squeezing was similar efficient as by endocytosis. This was confirmed by qPCR experiments, which indicated that about 80 % or 70 % of the total amount of **DON-1** were incorporated in the cellular fraction in the case of squeezing and endocytosis, respectively (Supplementary Figure S6). This value is similar to the delivery efficacy determined previously for siRNA, proteins, dextrane or carbon nanotubes^5^.

**Figure 2.**
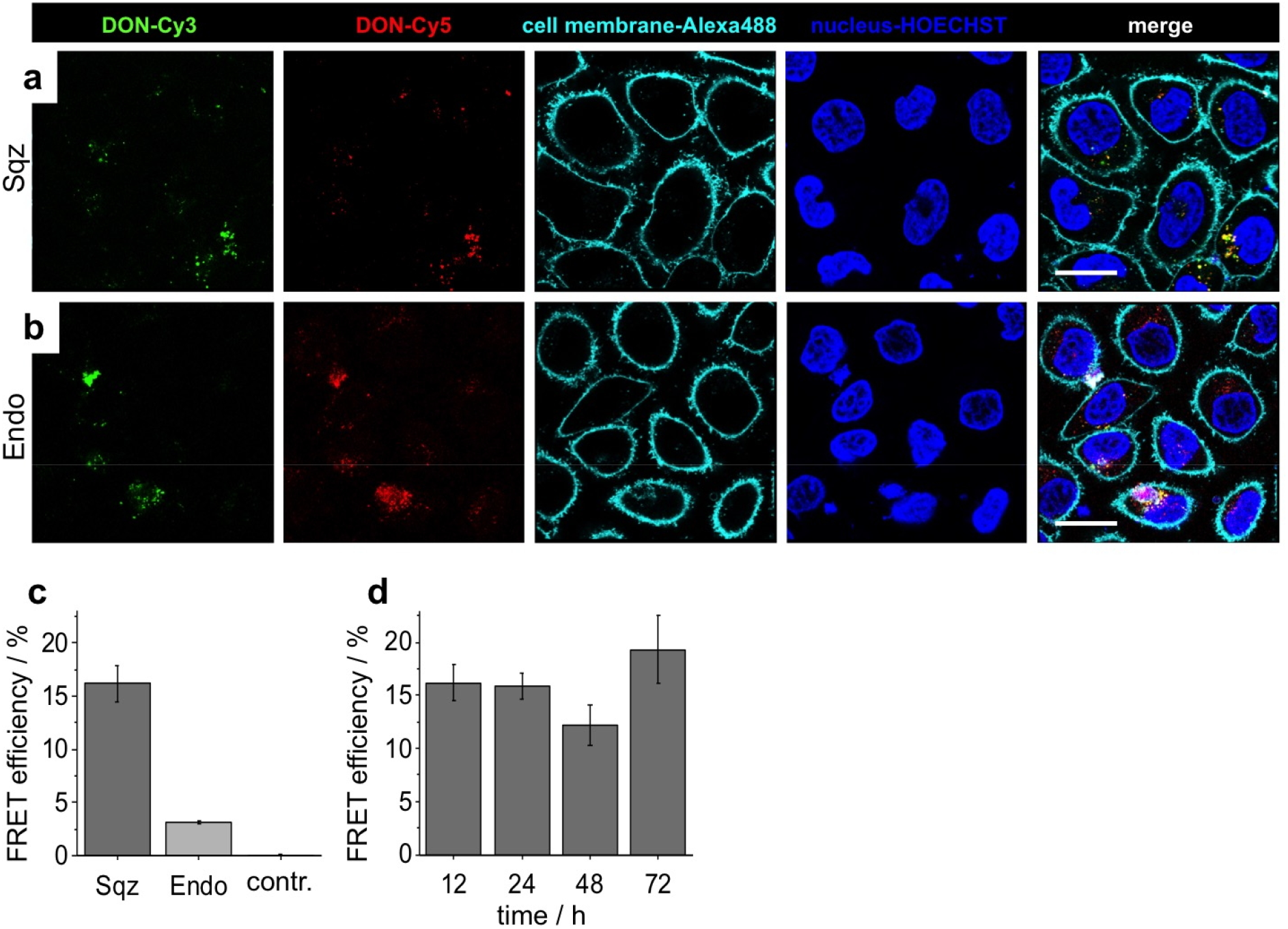
Cytosolic delivery of DON-1. Representative fluorescence micrographs of HeLa cells after delivery of **DON-1** by cell squeezing (**a**) or mere incubation leading to endocytosis (**b**). Cy3 and Cy5 fluorescence is visible in the green and red channels, respectively. The cell membrane was stained with wheat-germ agglutinin (WGA), conjugated to Alexa488 (cyan), cell nuclei were counterstained with HOECHST 33342 (blue). All images were taken 12 h after squeezing/endocytosis. Scale bar: 25 μm. (**c**) Quantification of FRET efficiency of **DON-1** in live HeLa cells, determined 12 h after delivery by squeezing (Sqz) or endocytosis (Endo). The control sample was heat-denatured (95 °C, 10 min) before squeezing. (d) FRET efficiency of **DON-1** determined at different time points after squeezing into HeLa cells.

Quantification of the FRET efficiency by using the acceptor photobleaching method revealed significant differences between DON samples delivered by squeezing or endocytosis. Indeed, FRET efficiency was significantly higher when **DON-1** was mechanically transfected as compared to the endocytic uptake (Figure 2c) and FRET signals remained constant over prolonged culturing periods (Figure 2d). Importantly, control experiments, wherein **DON-1** was denatured by heating to 95 °C prior to squeezing, led to no detectable FRET signals due to the disassembled Cy3/Cy5 FRET pairs (Figure 2c). The markedly reduced FRET of the endocytosed DON is presumably due to degradation of the DNA nanostructure inside lysosomal compartments, as suggested by colocalization experiments using a Cy5-labeled DON and stained lysosomes (Supplementary Figure S7). These results are in agreement with studies on the endocytosis of DNA nanostructures^40, 41^.

Quantification of cell viability confirmed the expectation that the squeezed cells are permeable immediately after the micromechanical treatment but almost fully recover after about 24 h (Supplementary Figure S8). In agreement with earlier studies that indicated the stability of DNA origami structures in cell culture medium and cell lysate for at least 24 h^21, 22^, our results confirmed that the supramolecular structure of **DON-1** remained intact and was not notably disturbed by either the mechanical transfection or intracellular components in the cytosol.

### Multiprotein-DON assemblies can be squeezed into cells

To investigate whether protein-decorated DONs can be squeezed into cells, we used **DON-2** that was loaded with 12 Cy3-labeled STV molecules. The resulting construct, **DON-2**-STV, was transfected into HeLa cells by either cell squeezing or endocytosis and the cells were analysed by confocal fluorescence microscopy. Clear signals of **DON-2-STV** could be derived from its Cy3 and Cy5 labels (green and red channels, respectively, in Figure 2a,b) only in the case of micromechanical transfection. These observations confirmed that the Cy3-labeled STV had passed the cell membrane aboard of the DON, thereby suggesting the intactness of the supramolecular structure of **DON-2-STV**. Surprisingly, the high uptake rate of STV-loaded **DON-2-STV** suggested also that the micromechanical delivery of protein-DNA constructs is even more effective than direct squeezing of proteins. In our hands, the squeezing of pure Cy3-labeled STV as well as of several other proteins was found to be inefficient (Table S3). For instance, repeated trials were necessary to successfully deliver the recombinant FLIP-HOB sensor protein into HeLa cells (Supplementary Figure S9). In contrast, no such problems were observed for proteins that were bound to the DNA nanostructures.

To investigate whether functional DNA nanopegboards can be delivered into the cytosol, we used construct **DON-3**, which contained to six chlorohexane (CH) binding sites in addition to six Cy5 labels (Figure 1d). The CH groups were used to selectively couple the self-ligating glucose sensor protein FLIP-HOB onto the DON and the resulting **DON-3-FLIP** construct was used for the micromechanical transfection of HeLa cells. Indeed, microscopy analysis of the transfected HeLa cells revealed the signals of the DON-attached Cy5 labels as well as of both fluorescent protein domains of the FLIP-HOB (eCFP, eYFP, Figure 3c). The functionality of the construct was verified by analysing the glucose-dependent eYFP/eCFP emission ratio change when the transfected HeLa cells were treated with medium containing either no glucose or 10 mM glucose (Figure 3c,d). It was evident by naked-eye inspection of the fluorescence micrographs that eYFP fluorescence was significantly reduced after replacing the 10 mM glucose medium with zero-glucose medium. Furthermore, quantitative analysis of small populations of about 10 cells within the visual field of the microscope, carried out for repeated changes of high- and low-glucose media, clearly indicated the reversibility of the I(527 nm)/I(476 nm) fluorescence ratio changes (Figure 3d). These results are indicative for the successful cytosolic delivery of functional multiprotein-DON constructs into living cells.

**Figure 3.**
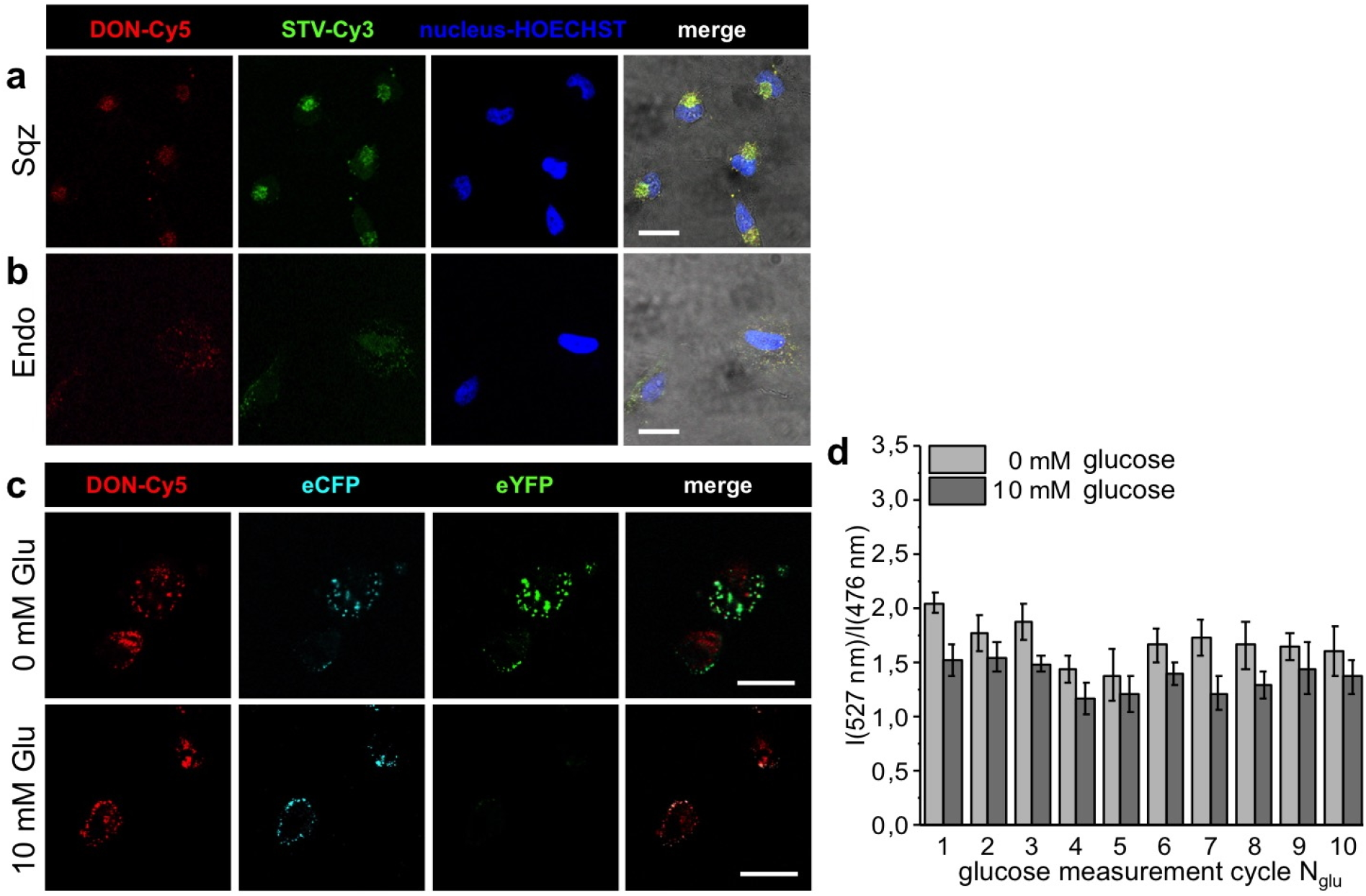
Cytosolic delivery of protein-decorated DON. Fluorescence micrographs of HeLa cells after delivery of **DON-2-STV** by cell squeezing or endocytosis, respectively (**a, b**). The cell nuclei were counterstained with HOECHST 33342 (blue). Scale bar is 25 μm. (**c**) Fluorescence micrographs of HeLa cells with **DON-3-FLIP** delivered by squeezing in medium with 0 und 10 mM glucose. Scale bar is 25 μm. (**d**) Quantitative analysis of the intensity ratio I(527 nm)/I(476 nm) in dependency of the glucose concentration and the measurement cycle N_glu_. In each cycle, the cells were exposed to medium with 0 mM (light gray bars) or 10 mM (dark gray) glucose and the corresponding intensity ratio was determined for the cells in the viewfield of the microscope.

### Cytosolic delivery and intracellular targeting of multiprotein-DON assemblies

We then investigated whether protein-decorated DONs can be targeted inside the cytosol of squeeze-transfected cells. To this end, we used construct **DON-2** that was initially loaded with STV and the DON-tethered proteins were then allowed to bind the biotinylated nuclear localization signal (NLS) peptide. The resulting construct, **DON-2-NLS**, was characterized by AFM (Figure 1b) and gel electrophoresis (Supplementary Figure S2) and used for transfection of HeLa cells by either cell squeezing or endocytosis. Fluorescence microscopy analysis (Figure 4a,b) clearly revealed that a co-localization of the **DON-2-NLS** with the cell’s nuclei occurred only when the construct was squeezed into the cells (Figure 4a), while no colocalization was observed for the endocytosed construct (Figure 4b). The results therefore indicate that the protein-decorated nanostructures were successfully delivered in the cytosol and specifically targeted to the nucleus.

**Figure 4.**
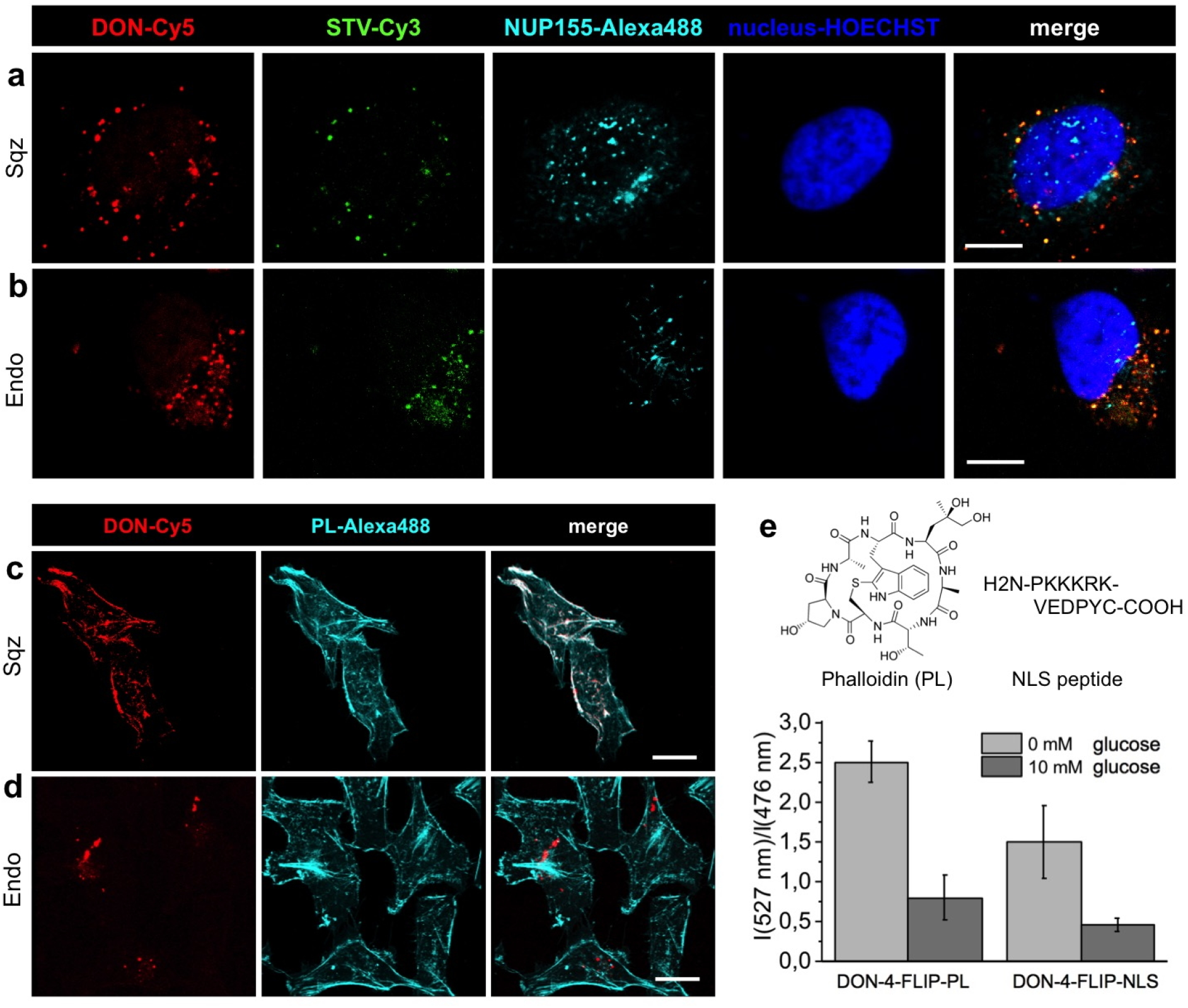
Cytosolic delivery and intracellular targeting of protein-functionalized DONs. Fluorescence micrographs of HeLa cells after delivery of **DON-2-NLS** by cell squeezing (**a**) or endocytosis (**b**). The nuclei and the nuclear pore complex component NUP155 of the fixed cells were counterstained with HOECHST 33342 (blue) and an Alexa488-labeled antibody directed against NUP155 (cyan), respectively. Scale bar is 10 μm. (**c, d**) Fluorescence micrographs of HeLa cells obtained after delivery of **DON-4-FLIP-PL** by squeezing (**c**) or endocytosis (**d**). Fixation and counterstaining was done with commercial PL-Alexa488. Scale bar is 25 μm. (**e**) Quantitative analysis of the intensity ratio I(527 nm)/I(476 nm) for intracellular **DON-4-FLIP-PL** and **DON-4-FLIP-NLS** in media with no or 10 mM glucose. Molecular structure and peptide sequence of PL and NLS are shown on top of the graph.

Next, we tested whether functional FLIP-decorated DONs can also be targeted to cytosolic compartments. To this end, we functionalised the **DON-4** construct with FLIP-HOB and STV proteins and the formed complex was then allowed to bind biotinylated PL. The resulting construct, **DON-4-FLIP-PL**, thus contained six copies of FLIP and four copies of PL-modified STV molecules in addition to six Cy5 labels (Figure 1e). The construct was then used for transfection of HeLa cells by squeezing or endocytosis. The cells were fixed after 5 h and the intracellular distribution of **DON-4-FLIP-PL** (red channel in Figures 4c, d) was determined by co-staining with commercial phalloidin-Alexa488 (cyan). Strikingly, the squeezed construct revealed a perfect co-localization with the filamentous actin, while the endocytosed construct showed intracellular aggregates. To verify the glucose biosensing capabilities of the targetable nanostructures, HeLa cells were transfected with either the **DON-4-FLIP-PL** or the **DON-4-FLIP-NLS** construct, both of which assembled as described above. Following to microfluidic squeezing, the cells were allowed to recover and then exposed to medium that contained either no or 10 mM glucose. Determination of the glucose-dependent eYFP/eCFP fluorescence of FLIP-HOB clearly indicated that both constructs were indeed capable of reporting the changes in the intracellular glucose level as the consequence of altered glucose concentrations in the cell culture media (Figure 4e).

## Discussion

We established a generic methodology for cytosolic delivery of large supramolecular complexes into living cells using DNA nanotechnology in combination with a micromechanical transfection platform that is fast, versatile in terms of transfected cargo and suitable for high-throughput campaigns^5, 30^. By using FLIP as a representative model for highly sensitive proteins that loose their functionality even upon slight perturbations of their tertiary structure and conformational dynamics, we confirmed that neither the immobilisation on DNA scaffolds nor the microfluidic delivery pose principal limitations to the here presented approach. Furthermore, the use of NLS peptide and Phalloidin as representative models for targeting groups clearly showed that our methodology has the potential to direct the delivered multiprotein assemblies to arbitrary locations inside living cells.

Further exploitation of our approach for the intracellular delivery of protein-decorated DNA nanostructures gives rise to a broad range of applications, in particular, for fundamental studies in cell biology. For instance, the cytosolic delivery of designed artificial multiprotein constructs will be useful to mimic, manipulate and interrogate naturally occurring supramolecular assemblies, such as the machinery involved in gene regulation, signal transduction, or cell division. A specific example concerns the reverse-engineering of partially assembled kinetochore complexes, a theme of increasing relevance for elucidating the architecture and functional dynamics of the human cell division machinery^6^. While seminal work on the DNA-directed assembly of tools for constitutional studies of large protein complexes had been restricted to the use of relatively simple DNA scaffolds for *in vitro* endpoint assays^42^, the now available methodology for cytosolic delivery of complex, multifunctional assemblies will open up a new chapter of this research.

More generally, the implementation of DNA nanotechnology allows for the use of an almost endless choice of functional protein complexes that is not limited to the here applied pseudo-2D origami plates but can be further extended to 3D DNA nanostructures, as exemplified by switchable hosts for protein encapsulation^25, 43^ Hence, the methodology described here will also be particularly useful for perturbation studies designed to unravel fundamentals of intracellular signalling^44^. DNA nanostructures have recently been established as powerful tools for biological perturbations at high spatiotemporal resolution, however, they are currently limited to receptor-mediated uptake of designed DNA nanostructures^29, 45^. Importantly, the micromechanical transfection allows to precisely tune the quantity of cargo for in-cell manipulation and this feature can be used to fine tune the biological output^30^.

More direct applications of our approach can be foreseen in the area of cellular immunotherapy. We observed that proteins tethered to DNA nanostructures are much better delivered into cells than pure proteins. Since this phenomenon seems counterintuitive due to the large difference in size, it calls for further systematic analysis of whether and how DNA and other nucleic acids can be used for improving the transfection efficiency of proteins. For instance, nucleic nanostructures could be specifically designed to boost the response of dendritic cells upon microfluidic transfection. Based on these examples, we believe that our approach will open up the door to novel tools and concepts to tackle questions in fundamental and applied biomedical research that cannot be answered by the established methodologies.

## Methods

### Assembly and purification of DNA origami nanostructures (DON)

The design of the unmodified rectangular DNA origami nanostructure (DON) assembled from the single stranded scaffold 109Z5 and 180 staple strand oligonucleotides (listed in Table S1) was carried out as previously described.^31^ The oligonucleotides were purchased from Sigma Aldrich. The CH-modified oligonucleotides were synthesized from the corresponding amino-modified oligonucleotides. All sequences and detailed information are provided as Supplementary Information. Analogous to Rothemund’s procedure^8^, 20 nM scaffold strand 109Z5 was mixed with 10-fold molar excess of staples in TEMg (20 mM Tris base, 2 mM EDTA, 12.5 mM MgCl_2_, pH 7.6 adjusted with HCl). Annealing was performed in a Thermocycler (Eppendorf Master cycler ® pro) by denaturing at 95 °C for 5 min and decreasing the temperature from 75 °C to 25 °C with a cooling rate of −0.1 °C/s. DON structures were purified by PEG-precipitation^46^ and the concentration was determined by qPCR^36^.

### DON decoration with proteins and biotin-modified targeting groups

20 μL 50 nM DON solution in TEMg were mixed with 1 to 4 equivalents FLIP-HOB per binding site (**DON-3, DON-4**) and/or 3 to 5 equivalents streptavidin (**DON-2, DON-4**) and incubated for 1 h (in case of further coupling wih additional biotinylated molecules) or 2 h at room temperature. 0.5 equivalents biotin-modified molecules NLS or PL per STV binding site were added to STV-modified DONs and incubated at room temperature for 1 h. Details on cloning of FLIP-HOB, protein expression and further characterization methods are given in the Supplementary Information.

### AFM analysis

Typically, samples were diluted up to 25 fold in TEMg (20 mM Tris base, 2 mM EDTA, 12.5 mM MgCl_2_, pH 7.6). 5 μl of this sample were deposited on a freshly cleaved mica surface (Plano GmbH) and adsorbed for 3 min at room temperature. After addition of 10 μl TAEMg (40 mM Tris, 20 mM acetic acid, 2 mM EDTA, 12.5 mM Mg acetate, pH 8.0), the sample was scanned with sharpened pyramidal tips (SNL-10 tips 0.35 N/m, Bruker) in Tapping Mode with a MultiModeTM 8 microscope (Bruker) equipped with a Nanoscope V controller. All images were analysed by the NanoScope software. Details on statistical AFM analysis are provided in the Supplementary Information.

### Confocal fluorescence microscopy and FRET measurements

Confocal fluorescence microscopy was performed on a Leica TCS-SP5 DMI 6000 (Leica). FRET was measured by using the FRET acceptor photobleaching method^47^ (Cy3/Cy5) or by determining the donor and acceptor intensity ratios (eCFP/eYFP of FLIP-HOB) with excitation at 436/20 nm in combination with a neutral density filter (1 % transmission) and two emission filters at 480/40 nm for eCFP and 535/30 nm for eYFP. Leica Application Suite Advanced Fluorescence (LAS AF) was used for image analysis and determination of the Pearson’s correlations coefficients. Further details on sample staining procedures are given in the Supplementary Information.

### Delivery of DONs by squeezing and endocytosis

For the micromechanical transfection of functionalized DON by squeezing, 1.10^5^ HeLa cells were suspended in 100 μL of a 10 nM DON solution in TEMg or PBS and were transferred to the squeezing device (SQZbiotech) combined with microfluidic chips (SQZbiotech) with channel dimensions of 10 × 7, 10 × 8 or 10 × 9 (length / μm × diameter / μm). The cells were pressed through the channels by applying a pressure of 20, 30, 40 or 50 psi. The cells were collected immediately and incubated on ice for 5 min. The DON solution was removed by centrifugation (800 g, 5 min) and the cell pellet was resuspended in 2 mL EMEM (Pan Biotech) or glucose free DMEM (Sigma Aldrich). The cell suspension was transferred to confocal dishes (μ-Slide 4 Well ibiTreat or μ-Dish 35 mm, high, ibiTreat: Ø 35 mm, high wall, Ibidi) and the cells were allowed to adhere at standard culturing conditions for at least 5 h. For the endocytosis controls, 1.10^5^ HeLa cells were suspended in 100 μL of 10 nM DON-solution, 2 mL medium was added and the cells were incubated for 12 h. Squeezing experiments with proteins were performed with the conditions listed in supplementary Table S3.

### Intracellular glucose assay

For the measurement of intracellular glucose sensor activity, the cells were treated with 2 mL glucose free DMEM und 10 mM glucose in DMEM with intermittent washing steps using DPBS (-/-) (Life Technologies). After each medium exchange, the fluorescence intensity values of eCFP and eYFP were recorded for about 10 cells by confocal microscopy.

## Acknowledgements

This work was supported in part by Deutsche Forschungsgemeinschaft in the course of priority programme SPP 1623 and by the Helmholtz programme BioInterfaces in Technology and Medicine. We thank Anke Dech and Cornelia Ziegler for experimental help and Dr. Parvesh Wadhwani of KIT’s IBG-2 for the synthesis of the NLS peptide.

## Author contributions

P.M.N. and A.S. designed and performed the cell squeezing experiments. K.J.K, A.A., and J.B. designed, synthesized and characterized DON complexes. K.S.R. and K.J.K. designed and cloned recombinant fusion proteins. R.T. and A.K. established design and implementation of initial squeezing experiments. K.J.K. and C.M.N. analysed the data and wrote the manuscript. C.M.N. designed the project and directed the work. All authors discussed the results and commented on the manuscript.

